# Permeabilization-free *en bloc* immunohistochemistry for correlative microscopy

**DOI:** 10.1101/2020.10.07.327734

**Authors:** Kara A Fulton, Kevin L Briggman

## Abstract

A dense reconstruction of neuronal synaptic connectivity typically requires high-resolution 3D electron microscopy (EM) data, but EM data alone lacks functional information about neurons and synapses. One approach to augment structural EM datasets is with the fluorescent immunohistochemical (IHC) localization of functionally relevant proteins. We describe a protocol that obviates the requirement of tissue permeabilization in thick tissue sections, a major impediment for correlative pre-embedding IHC and EM. We demonstrate the permeabilization-free labeling of neuronal cell types, intracellular enzymes, and synaptic proteins in tissue sections hundreds of microns thick in multiple brain regions while simultaneously retaining the ultrastructural integrity of the tissue. Finally, we explore the utility of this protocol by performing proof-of-principle correlative experiments combining two-photon imaging of protein distributions and 3D electron microscopy.

## Introduction

Anatomical reconstructions of synaptic connectivity are one essential component of understanding neuronal circuits in the nervous system. Recent improvements in automating the collection of 3D electron microscopy (EM) data have dramatically increased the tissue volumes that can be acquired (Briggman and Bock, 2012; Eberle et al., 2015). Connectivity data derived from EM, however, does not alone contain sufficient information about the functional properties of neurons to fully constrain biologically plausible computational models. One approach to augment EM datasets with functional information is to physiologically record from neurons in a tissue sample prior to fixation (Bock et al., 2011; Briggman et al., 2011; Lee et al., 2016; Wanner and Friedrich, 2020). As a complementary approach, fluorescence immunohistochemistry (IHC) can be used to identify the constituent proteins of neuronal cell types and synapses that are key determinants of the functional properties of neurons (Coons et al., 1942; Cuello et al., 1983).

Post-sectioning IHC is typically employed for correlative EM because the sectioning process provides access to intracellular epitopes. For example, array tomography was developed to multiplex the labeling of numerous targets in ultrathin tissue sections (Collman et al., 2015; Micheva and Smith, 2007). However, this approach is incompatible with *en bloc*-based methods for collecting 3D EM data that require pre-embedding IHC, as in the case of focused ion beam scanning electron microscopy (FIB-SEM, (Hayworth et al., 2015; Knott et al., 2008)), gas cluster ion beam SEM (Hayworth et al., 2020), and serial block-face scanning electron microscopy (SBEM, (Denk and Horstmann, 2004)). For pre-embedding IHC, a membrane permeabilization step with nonionic surfactants such as Triton X-100 or polysorbate 20 (Tween 20) is typically performed to enable antibody penetration into thick tissue blocks, but at the cost of degraded tissue ultrastructure (Helenius and Simons, 1975; Humbel Bruno et al., 1998).

To address this limitation, we began with the observation that the preservation of extracellular space (ECS) during chemical tissue fixation improves the penetration of antibodies into the mammalian retina using minimal permeabilization (Pallotto et al., 2015). Building on this finding, we developed a protocol that is capable of immunohistochemically labeling proteins throughout brain tissue sections up to one millimeter thick without permeabilization. Here we describe the key changes that were made in comparison to conventional IHC protocols leading to improved ultrastructural preservation. In addition, we demonstrate several applications of the protocol including the labeling of cell-type specific proteins in various brain regions and proof-of-principle correlative light microscopy (LM)/EM experiments.

## Results

### Optimization of a permeabilization-free protocol to label ECS preserved tissue sections

Aldehyde-based fixation of the brain has been previously demonstrated to result in the loss of the 15 - 20% ECS volume fraction normally found *in vivo* (Van Harreveld and Malhotra, 1967). The observation that detergent permeabilization is required for antibodies to penetrate into thick brain tissue sections has largely been based on such, usually perfusion-fixed, tissue in which ECS is not preserved. We began by demonstrating this requirement using a fluorophore-conjugated primary antibody for the neuronal soma marker, NeuN, as a representative antibody (Mullen et al., 1992) and prepared 300 μm thick perfusion-fixed sections of mouse cerebral cortex. When detergent was omitted from the labeling protocol, we observed a gradient of fluorescently labelled somata that decreased in the center of sections, as expected, and the ultrastructural membrane integrity of the tissue remained intact (Figure 1b). With the inclusion of a commonly used detergent, Triton X-100, neuronal somata were uniformly labeled throughout the depth of the sections but at the cost of severely degraded membrane integrity due to lipid solubilization (Figure 1a).

**Figure 1.**
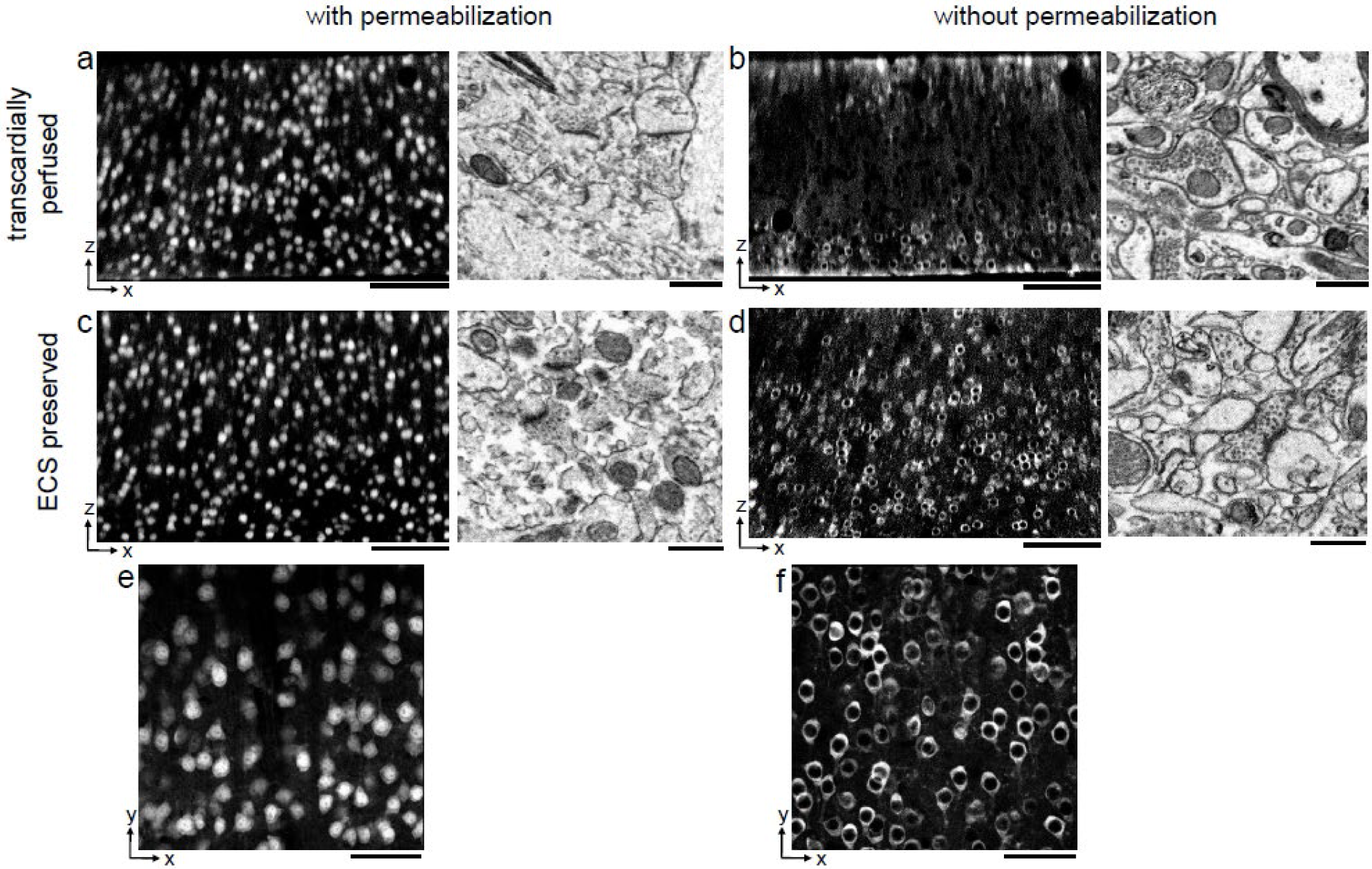
Permeabilization-free labeling in ECS preserved cerebral cortex. **a,b)** 300 μm thick sections from the cerebral cortex of a transcardially perfused mouse. Incubation with Alexa Fluor-488 conjugated anti-NeuN was performed with (a) or without (b) 0.3% Triton to label neuronal somata. X-Z reslices of 2P image volumes (left panels) are 10 μm average intensity projections from the center of the sections. Sections were then stained for EM and a region from the center of the section was examined for ultrastructural integrity (right panels). **c,d)** Same procedure as in (a,b) but for 300 μm thick acute ECS preserved sections from the cerebral cortex of a mouse. **e,f)** 10 μm average intensity projections of X-Y slices from center of the image volumes highlighting nuclear exclusion of anti-NeuN labeling when Triton is omitted (f). Scale bars: a-d, left panels: 100 μm; a-d, right panels: 1 μm; e,f: 50 μm.

We then replicated this experiment in 300 μm thick acute immersion-fixed mouse cortical sections in which ECS was preserved as recently described (Pallotto et al., 2015). As with perfusion-fixed sections, the inclusion of a detergent in the protocol yielded uniform penetration of anti-NeuN at the expense of degraded membrane integrity (Figure 1c). However, when detergent was omitted from the protocol, the labeling of neuronal somata in ECS-preserved sections remained uniform throughout the tissue (Figure 1d).

Importantly, the exclusion of a permeabilization step yielded intact lipid membranes based on electron micrographs from the same tissue (Figure 1d). This result was repeatable between cortical sections and across individual mice (Suppl Figure 1). We noted a nuclear exclusion of the NeuN labeling in the non-permeabilized sections (Figure 1f) compared to permeabilized sections (Figure 1e) indicating nuclear membranes remained impermeant to the antibody.

Based on the observation that anti-NeuN can penetrate thick ECS-preserved tissue sections without permeabilization, we proceeded to optimize the parameters of the protocol and explore the boundary conditions of the labeling uniformity. The parameters we titrated included fixative composition and duration, antibody concentration and duration of incubation, IHC buffer composition and osmolarity, incubation temperature, and the use of a tissue clearing protocol. The resulting optimized IHC protocol (Table 1) incorporates these systematic optimization steps. A balance between ultrastructural integrity and penetration depth was dependent on the concentrations of paraformaldehyde (PFA) and glutaraldehyde (GA) in the primary fixative solution, as expected based on the different cross-linking properties of the two aldehydes (Hopwood, 1985). A mixture of 4% PFA and 0.005% GA yielded a good compromise between penetration depth and EM quality, although small variations in the GA concentration led to similar results (Suppl Figure 2). A second key parameter was the concentration of the primary antibody. Uniform labeling was achieved when using IgG primary antibody concentrations in the range of 33 – 66 nM (Suppl Figure 3a). Additional key parameters were the use of an antibody incubation buffer that was isotonic with the fixation buffer, the ratio of NaCl to PB in the antibody incubation buffer, and room temperature processing (data not shown). Of the parameters we varied, the labeling uniformity was less sensitive to the duration of the initial fixation (Suppl Figure 3b) and the omission of blocking serums (Suppl Figure 4), the utility of which has been questioned (Buchwalow et al., 2011). The ultrastructural quality was less sensitive to the duration of the initial fixation (Suppl Figure 3b) and the duration of the antibody incubations (Suppl Figure 5).

**Table 1.** Optimized permeabilization-free IHC protocol. Incubation solutions, durations and temperatures are reported along with reference to relevant supplementary figures. The duration of antibody incubation was varied dependent on section thickness. Asterisk (*) indicates PB concentration should be varied to achieve desired ECS volume fraction in a tissue dependent manner. Abbreviations: artificial cerebral spinal fluid (ACSF), paraformaldehyde (PFA), glutaraldehyde (GA), sodium phosphate buffer (PB), antibody (Ab), sodium cacodylate buffer (CB), NIRB (near-infrared branding), RT (room temperature), overnight (o/n).

| <u>Solution</u> | <u>Duration</u> | <u>Temp.</u> |  | <u>Relevant Supplemental Figures</u> |
| --- | --- | --- | --- | --- |
| ACSF | ~10 min | 4° C | Dissection,<br>acute sectioning |  |
| 4% PFA + 0.005% GA<br>in 0.175 M PB*, pH 7.4 | 4 - 16 hr | 4° C | First fixation | Suppl Fig 2 - Optimization of fixation parameters<br>Suppl Fig 3 - Optimal duration of primary fixation |
| 0.05 M glycine in<br>0.175 M PB, pH 7.4 | o/n | RT | Glycine rinse |  |
| 0.175 M PB, pH 7.4 | 4 hr | RT | Rinse |  |
| 0.17 M NaCl +<br>0.01 M PB + 1° Ab | 2 - 7 days | RT | Primary antibody | Suppl Fig 4 - Effect of serum blocking<br>Suppl Table 2 - Antibodies tested |
| 0.175 M PB, pH 7.4 | o/n | RT | Rinse |  |
| 0.17 M NaCl +<br>0.01 M PB + 2° Ab | 2 - 7 days | RT | Secondary antibody | Suppl Fig 5 - Effect of prolonged antibody incubation |
| 0.175 M PB, pH 7.4 | 4 hr - o/n | RT | Rinse |  |
| 2% PFA in 0.175 M PB,<br>pH 7.4 | 2 hr | RT | Second fixation |  |
| 0.05 M glycine in<br>0.175 M PB, pH 7.4 | 4 hr | RT | Glycine rinse |  |
| 0.175 M PB, pH 7.4 | 4 hr | RT | Rinse |  |
| 20 → 100% fructose | 20 - 48 hr | RT | SeeDB refractive index<br>matching | Suppl Fig 6 - Effect of SeeDB clearing |
| 100% wt/vol or<br>80.2% wt/wt fructose | < 1 day | RT | 2P imaging & NIRB |  |
| 0.175 M PB, pH 7.4 | 12 hr | RT | Return tissue to PB |  |
| 0.15 M CB, pH 7.4 | 2 hr | 4° C | Rinse |  |
| 2% GA in 0.15 M CB,<br>pH 7.4 | 2 hr | 4° C | Third fixation |  |
| 0.15 M CB, pH 7.4 | o/n | 4° C | Rinse |  |
|  | 2 days |  | EM staining/embedding |  |
|  |  |  | EM imaging |  |

Because we prepared relatively thick fixed tissue sections, we were unable to image fluorescence throughout the depth of the sections and therefore employed a tissue clearing method. Many of the tissue clearing protocols recently described are not compatible with correlative LM/EM due to a degradation of tissue ultrastructure (Richardson and Lichtman, 2015). We therefore chose to use a refractive index matching approach utilizing high concentration fructose solutions, SeeDB, that does not involve the dissolution of membrane lipid (Ke et al., 2013). We confirmed that re-fixing the tissue with 2% PFA before serially incubating in fructose solutions retains ultrastructural quality (Suppl Figure 6). The use of SeeDB clearing allowed us to obtain two-photon (2P) excited fluorescence signals throughout the depth of 300 μm fixed tissue sections (Denk et al., 1990).

To explore the upper bound of the depth penetration achievable with the permeabilization-free protocol, we prepared 1 mm thick mouse cortical coronal sections and varied the fixative buffer concentration to yield different ECS volume fractions as previously described (Pallotto et al., 2015) (Figure 2). The duration of the anti-NeuN incubation was increased from 72 hrs (for 300 μm sections) to 120 hrs to allow sufficient time for penetration. We observed that the depth of penetration increased as the fixative buffer concentration increased and that the highest concentration we used yielded complete penetration through 1 mm thick sections (Figure 2c). As a result of the prolonged incubation duration, somatic nuclei near the edge of the section were occasionally labeled whereas those nuclei in the center exhibited exclusion of the antibody indicating a time dependence of the nuclear exclusion.

**Figure 2.**
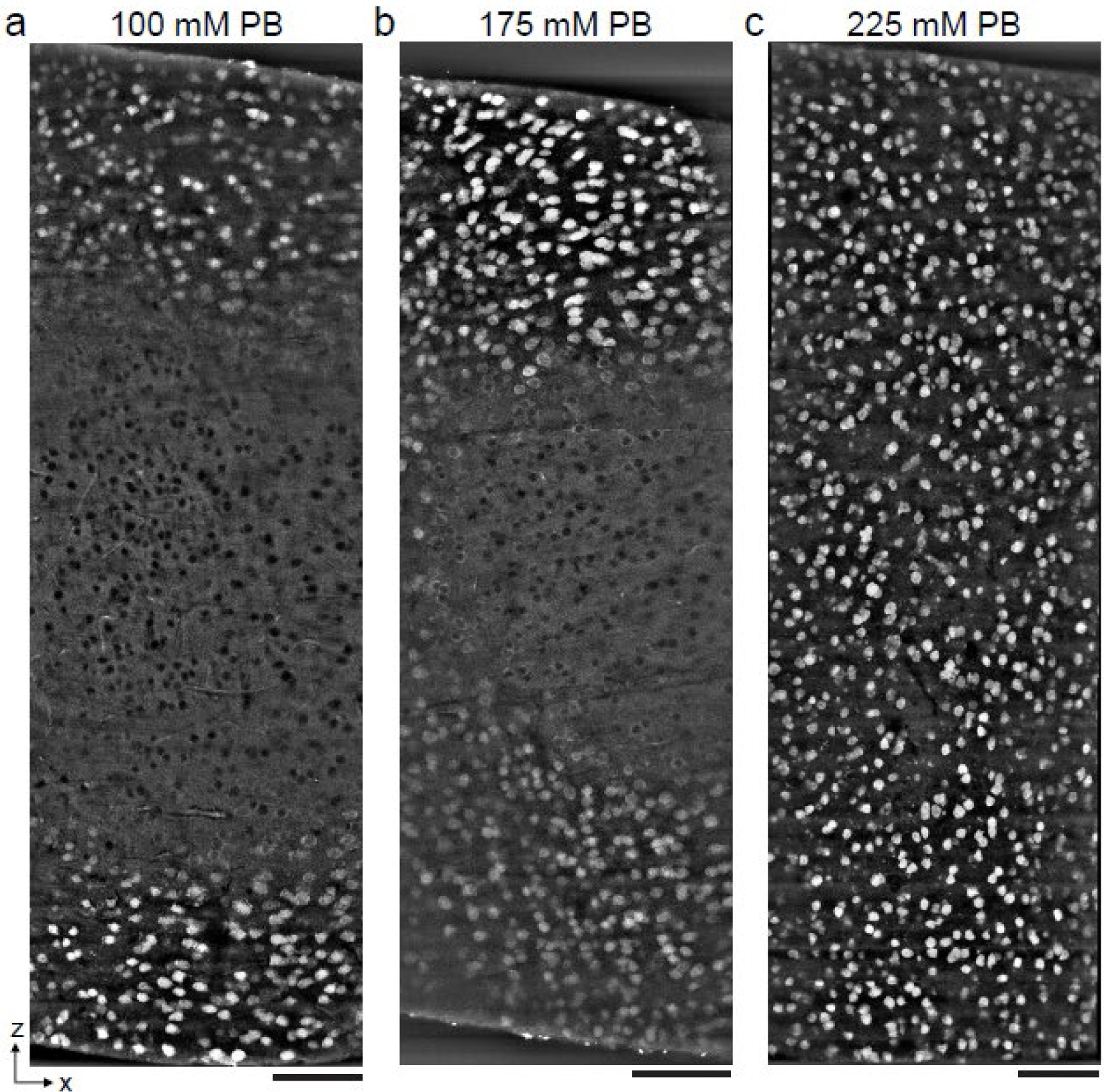
Antibody depth penetration increases with fixation osmolarity. Approximately 4 × 1 × 1 mm thick acute ECS preserved sections from the mouse cerebral cortex, fixed with increasing buffer concentrations to yield a) low (215 mOsm), b) medium (360 mOsm), or c) high (480 mOsm) osmolarities corresponding to increasing ECS volume fractions. Incubation with Alexa Fluor-488 conjugated anti-NeuN was performed without Triton to label neuronal somata. Tissue sections were hemisected and the cut surface was 2P imaged to investigate the penetration depth. Images are 10 μm maximum intensity projections from the cut surface. Scale bars: 100 μm.

### Distinguishing neuronal cell types and synaptic proteins with permeabilization-free IHC

During our optimization of the protocol parameters, we used anti-NeuN as a general label of neuronal somata. We next explored whether the protocol is compatible with additional antibodies and whether we could multiplex the immunofluorescent labeling of multiple protein targets. We first compared the labeling of neurons expressing the calcium-binding proteins calretinin (CR) and calbindin (CB) in the mouse cerebral cortex (Figure 3a,b). The CR+ and CB+ cell densities labeled with the permeabilization-free protocol (Figure 3b) were similar to those labeled when a permeabilization step was included (Figure 3a). Similar to the results with anti-NeuN, the permeabilization-free protocol generally yielded nuclear exclusion of the CR and CB labeling (Figure 3b).

**Figure 3.**
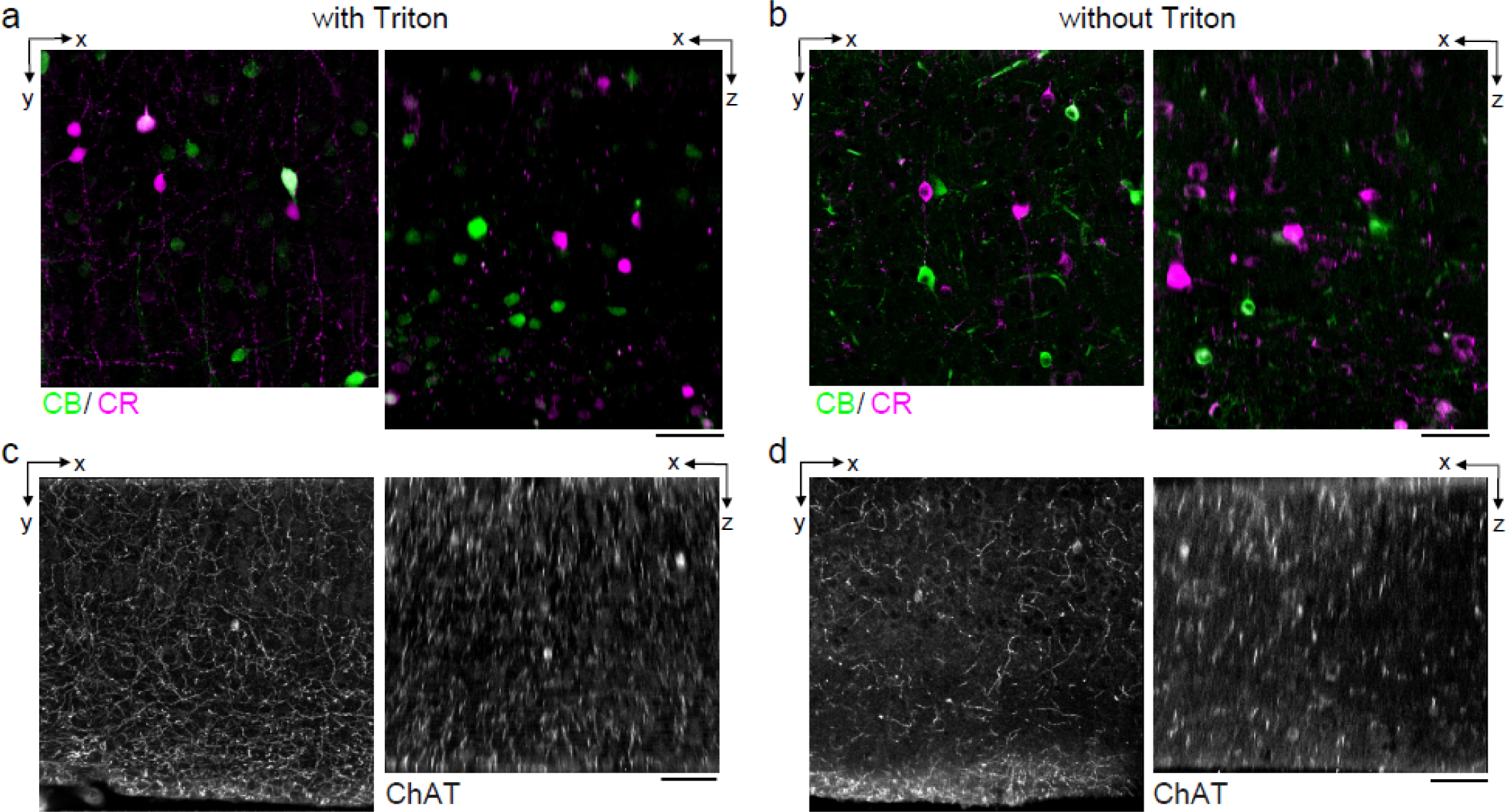
Permeabilization-free labeling of cell types and axons. **a,b)** 300 μm thick acute ECS preserved sections from the mouse cerebral cortex. Simultaneous incubation with anti-calbindin (CB) and anti-calretinin (CR) was performed with (a) and without (b) 0.3% Triton to label interneuron somata. X-Y slices (left panels) and X-Z reslices (right panels) of 2P image volumes are 10 μm average intensity projections from the center of the sections. **c,d)** Same as in (a,b) but 300 μm thick acute ECS preserved sections from the mouse medial prefrontal cortex were labeled with anti-choline acetyl transferase (ChAT) with (c) and without (d) 0.3% Triton to label cholinergic axons. Scale bars: 50 μm.

We next examined the IHC labelling of the cytosolic enzyme choline acetyltransferase (ChAT) expressed in cholinergic neurons. We were motivated to explore whether cholinergic axons could be labeled in brain regions distant from their respective somata. Using tissue sections from the medial prefrontal cortex (mPFC), we achieved labeling of ChAT+ axons both with and without permeabilization (Figure 3c,d) consistent with previous reports of the innervation patterns in the mPFC (Zhang et al., 2010). However, we noted a discontinuity of the axonal labeling in the permeabilization-free sections (Figure 3d) compared to the permeabilized sections (Figure 3c). We hypothesize that the incomplete axonal labeling may be due to a narrowing of the diameter of portions of the axons and therefore present a hindrance to the diffusion of the antibodies or, potentially, that myelination may limit penetration into axons. Both of these possibilities are consistent with the lipid solubilizing effect of permeabilization that would improve penetration into axons, but at the cost of membrane integrity.

We then explored whether synaptic proteins in the mouse cortex could be labeled without permeabilization. We first attempted to label the postsynaptic density protein 95 (PSD-95), but found poor labeling without permeabilization (data not shown). Among postsynaptic proteins, super-resolution microscopy has demonstrated that PSD-95 is within approximately 30 nm of the postsynaptic membrane (Dani et al., 2010). We hypothesized that, in the absence of solubilizing lipids with Triton, there may be a steric hindrance for antibodies to label proteins that are in such close proximity to the plasma membrane. We therefore attempted to label Homer, a protein found more distant, greater than 50 nm, from the postsynaptic membrane of excitatory synapses (Dani et al., 2010). The labeling pattern of Homer (Figure 4b) was similar to permeabilized sections (Figure 4a) throughout the depth of 300 μm thick sections. We noted a larger variability in the fluorescence intensity of labeled puncta in the non-permeabilized tissue compared to permeabilized tissue.

**Figure 4.**
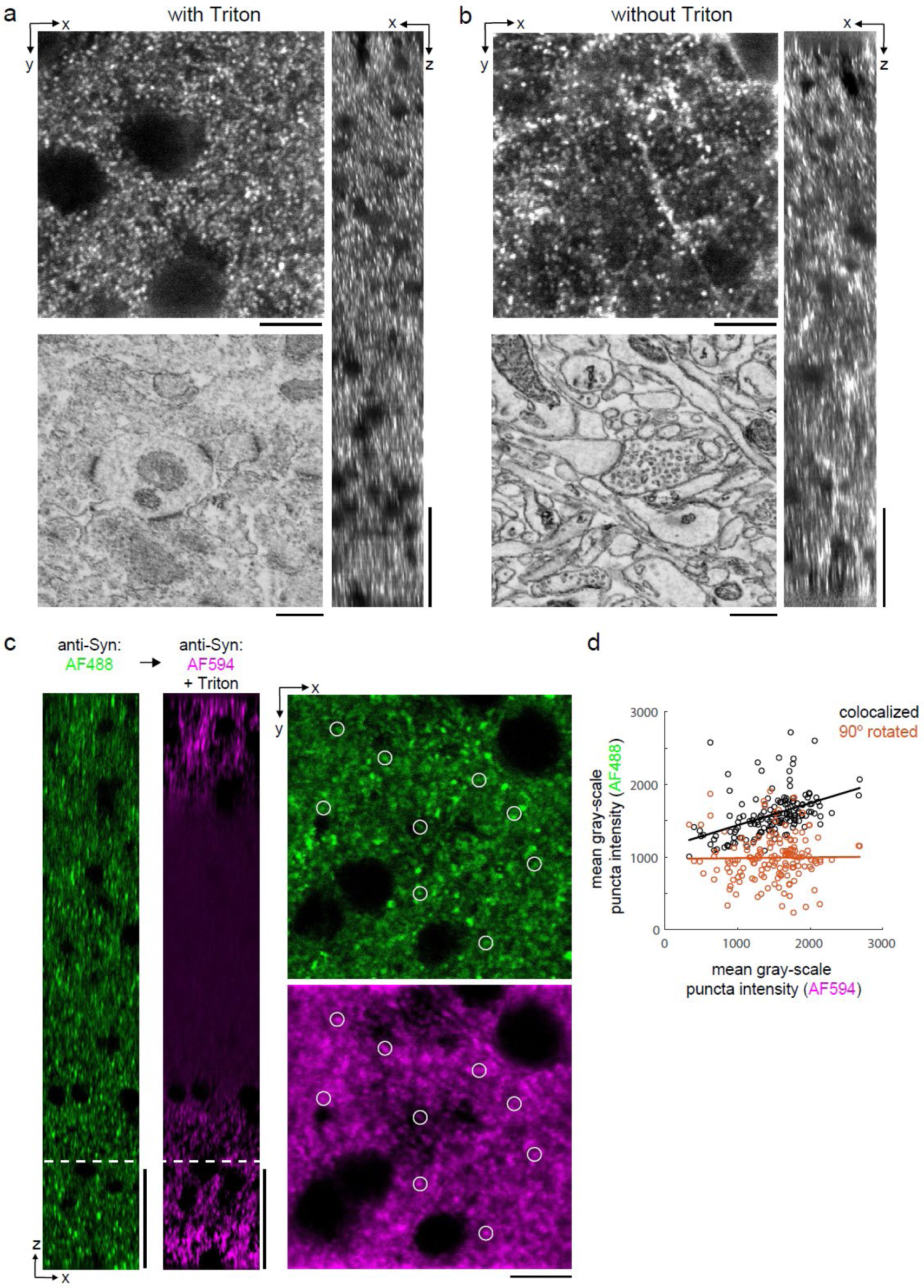
Permeabilization-free labeling of synaptic proteins. **a,b)** 300 μm thick acute ECS preserved sections from the mouse cerebral cortex. Incubation with anti-Homer was performed with (a) and without (b) 0.3% Triton. X-Y slices (upper left panels) and X-Z reslices (right panels) of 2P image volumes are taken from the center of the sections. Sections were then stained for EM and a region from the center of the section was examined for ultrastructural integrity (lower left panels). **c)** 300 μm thick acute ECS preserved slices from the mouse cerebral cortex. Sequential incubation with Alexa Fluor-488 (AF488) conjugated anti-synaptophysin (anti-Syn) without Triton (left, green) followed by Alexa Fluor-594 (AF594) conjugated anti-Syn with 0.3% Triton (left, magenta) shown as X-Z reslices. Puncta were outlined in X-Y images at the depth indicated (left, white dashed line) in the AF594 channel (lower right panel, representative white circles) for co-localization analysis with the AF488 channel (upper right panel). **d)** Co-localization analysis of 150 puncta from image stacks in (c) comparing co-localized puncta (black) to a 90 degree rotation of the AF488 channel (orange). Scale bars: a-b (upper left panels: 10 μm; right panels: 50 μm; lower left panels: 0.5 μm), c (left panels: 50 μm; right panels: 10 μm).

Finally, we explicitly compared the labeling efficiency of a presynaptic protein, synaptophysin, in the same tissue section before and after permeabilization. We took advantage of the availability of synaptophysin antibodies directly conjugated to Alexa Fluor (AF) molecules of different excitation wavelengths (AF488 and AF594). We first labeled mouse cortical sections with AF488 anti-synatophysin without permeabilization and then labeled the same sections with AF594 anti-synatophysin including Triton permeabilization. Prior to permeabilization, we observed a labeling density consistent with published labeling patterns (Figure 4c, green) (Grant et al., 2016). Following permeabilization, the labeling pattern was similar at the edges of the tissue but the antibody labeling was notably absent in the center of the section (Figure 4c, magenta). This result is consistent with the observation that treatment with Triton X-100 can lead to the extraction of synaptophysin from fixed tissue (Hannah et al., 1998). To distinguish whether any presynaptic puncta were labeled following permeabilization that were not labeled prior to permeabilization, we quantified the co-localization of puncta in the two fluorescence channels (Figure 4d). Puncta intensities (n = 150 puncta) were significantly correlated (linear regression t-test, p = 1.5 × 10^−8^) compared to a 90 degree rotation of the images (p=0.86), indicating a co-localization of the labeling of synaptophysin pre- and post-permeabilization

### Applicability of permeabilization-free labeling across different brain regions

In addition to the mouse cerebral cortex, we assessed antibody penetration in brain regions with different cytoarchitectures including the mouse hippocampus and olfactory bulb. Labeling 300 μm thick tissue sections from these brain regions with antibodies targeting NeuN, CR, and CB resulted in patterns similar to permeabilized sections (Figure 5). In the hippocampus, we observed the CR+ subgranular band and granular cell layer positioning of CB+ neurons as previously reported (Ohira et al., 2010) (Figure 5a-d). In the olfactory bulb, non-overlapping populations of CB+ and CR+ periglomerular cells (Panzanelli et al., 2007) were positively identified (Figure 5e-h).

**Figure 5.**
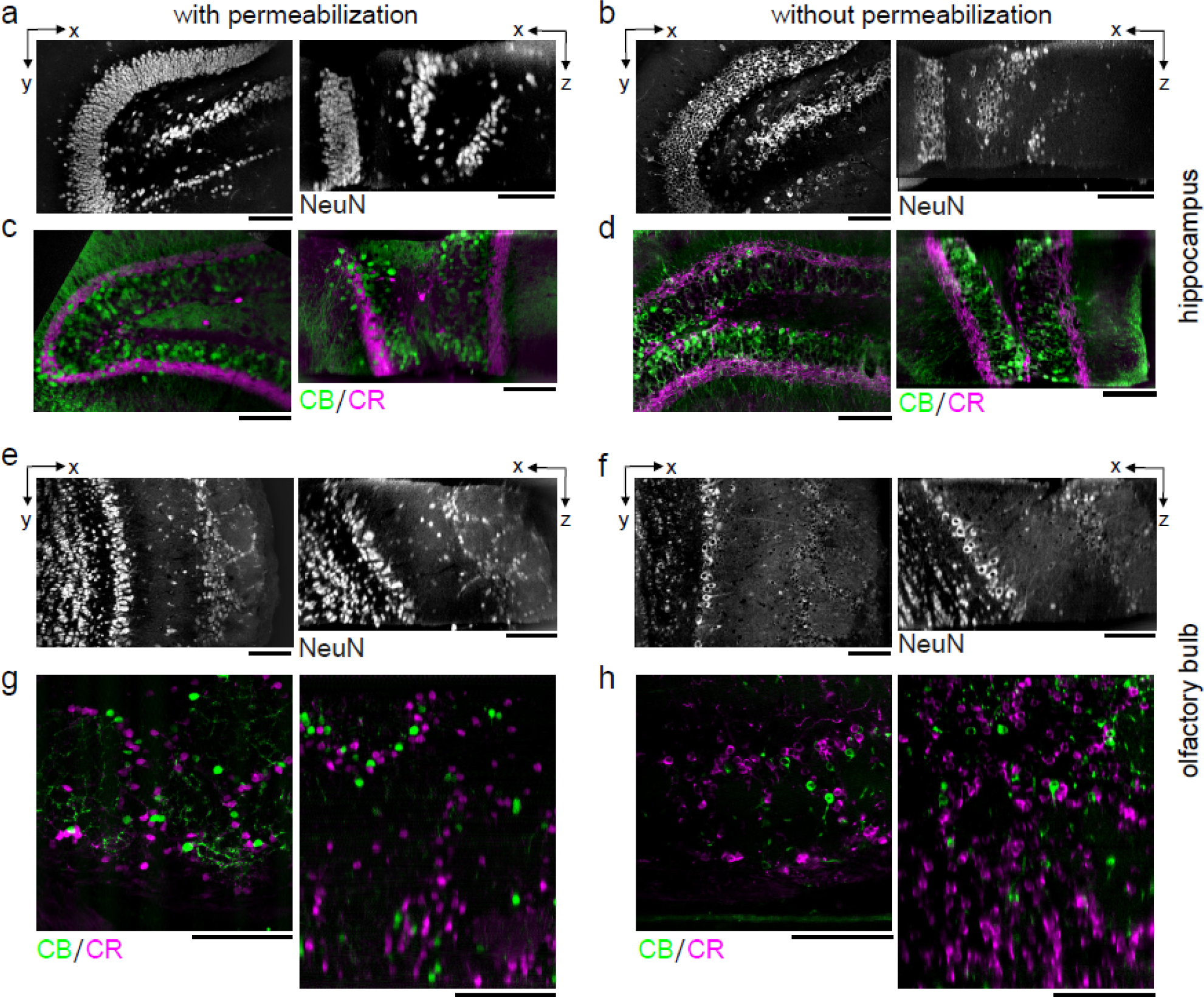
Permeabilization-free labeling of diverse brain regions. **a-d)** 300 μm thick acute ECS preserved sections from the mouse hippocampus. Incubation with Alexa Fluor-488 conjugated anti-NeuN (a,b) or simultaneous incubation with anti-calbindin (CB) and anti-calretinin (CR) (c,d) was performed with (a,c) and without (b,d) 0.3% Triton to label interneuron somata. X-Y slices (left panels) and X-Z reslices (right panels) of 2P image volumes are 10 μm average intensity projections from the center of the sections. **e-h)** 300 μm thick acute ECS preserved sections from the mouse olfactory bulb. Incubation with Alexa Fluor-488 conjugated anti-NeuN (e,f) or simultaneous incubation with anti-calbindin (CB) and anti-calretinin (CR) (g,h) was performed with (e,g) and without (f,h) 0.3% Triton to label neuronal somata. 2P image slices as in a-d. Scale bars: 100 μm.

### Permeabilization-free labeling for correlative LM/EM

Our primary motivation for developing a permeabilization-free IHC protocol was to enable the correlation of fluorescently labeled proteins with their location in EM images. As a first proof-of-principle, we investigated the application of converting GFP in labeled neurons into an electron dense signal. We retrogradely labeled mitral and tufted cells in the mouse olfactory bulb by injecting an adeno-associated virus (AAV) encoding Flex-eGFP in the mouse piriform cortex of a Pcdh21-Cre mouse (Wachowiak et al., 2013). Following 2P imaging of 300 μm thick acute sections (Figure 6a), the GFP was labelled with a primary anti-GFP antibody and a secondary antibody conjugated to horseradish peroxidase (HRP). The HRP-assisted polymerization of diaminobenzidine (DAB) was then carried out in the presence of hydrogen peroxide, yielding a DAB polymer that was strongly osmiophilic in EM images. Individual mitral and tufted cell neurites were readily distinguished from unlabeled neurites in the surrounding neuropil of the external plexiform layer of the olfactory bulb (Figure 6b-c).

Ultimately, the correlation of LM and EM is most informative when performed in 3D volumes. We therefore combined the permeabilization-free IHC protocol with a block-face EM technique as a correlative pipeline consisting of five steps (Table 1): 1) permeabilization-free *en bloc* IHC, 2) tissue clearing by refractive index matching (Ke et al., 2013), 3) 2P imaging of the tissue volume and near-infrared branding (Bishop et al., 2011) of a region of interest, 4) reversal of the tissue clearing protocol back to buffer, and 5) EM staining and acquisition of a SBEM volume.

As a second proof-of-principle, we labeled tyrosine hydroxylase (TH) positive axons in a 300 μm section from the mPFC (Zhang et al., 2010) (Figure 6d). Following SeeDB clearing, 2P imaging and branding of the section, we stained the tissue for EM and collected an 89 × 83 × 75 μm^3^ SBEM volume centered on the branded region (Figure 6e). The 2P and EM datasets were aligned by fitting an affine transform using landmarks in the two datasets including somata and blood vessels. We then locally searched regions of the EM dataset to identify the matching trajectories of fluorescent axons to those in the EM volume (Figure 6f,g) and identified TH+ axons within the EM volume (Figure 6h).

## Discussion

We have developed a protocol to immunohistochemically label thick tissue sections that omits the commonly used permeabilization step and is therefore compatible with correlative *en bloc* volume EM techniques. The method depends on the preservation of ECS during tissue fixation (Figures 1 & 2) and we suggest that the simplest explanation for the improved labeling is that the diffusion spaces that are preserved allow antibodies to diffuse deep into tissue and travel to close proximity of their antigens. That is, rather than translocating across multiple plasma membranes in densely packed neuropil, an antibody in ECS preserved tissue could, in principle, only need to cross one membrane to reach an epitope. The mechanism by which an antibody crosses a membrane that has not been permeabilized by a detergent is not clear to us, but we note a few observations. First, aldehyde fixation alone has been described to semi-permeabilize membranes due to their denaturizing effects on proteins (Hopwood, 1985). Second, the hydrodynamic radius of an IgG antibody is approximately 5 nm (Armstrong et al., 2004; Jøssang et al., 1988), meaning a small pore in a membrane of similar size is required to provide access to intracellular epitopes. We cannot rule out that the ECS preservation process itself leads to damage of some membranes allowing an antibody to enter through a damaged region and then diffuse along the intracellular cytosol of neurons. However, if such damage occurs, it has not prevented us from reconstructing neuronal morphologies and identifying synapses in ECS preserved tissue ((Pallotto et al., 2015); Figures 4 & 6). Applying the method to high-pressure frozen tissue in which ultrastructure and ECS is preserved in a more native state may yield additional insights into the mechanism (Korogod et al., 2015).

A current limitation of the protocol is our inability to label antigens that are closely associated with lipid membranes such as the postsynaptic protein PSD-95. The postsynaptic terminal is densely packed with protein and our ability to label the Homer protein, which is more distant from the postsynaptic membrane than PSD-95 (Dani et al., 2010), suggests a steric hindrance for antibody binding when a protein is in close proximity to the plasma membrane. In contrast, we were able to label the presynaptic protein synaptophysin located on the surfaces of vesicles that are relatively accessible within presynaptic terminals. Therefore, some degree of permeabilization may be a requirement for proteins that are located in close spatial proximity to the plasma membrane. One alternative would be to explore weaker detergents than Triton, such as low concentrations of Tween, which was previously used to label ECS preserved retinal tissue with satisfactory ultrastructure (Pallotto et al., 2015). Alternatively, if the size of IgG antibodies is limiting the binding to postsynaptic targets, the use of physically smaller antibodies, such as nanobodies (Hamers-Casterman et al., 1993), may improve labeling in combination with our protocol. Nanobodies have recently been shown to label targets up to 100 μm, but not deeper, from one surface of a tissue section (Fang et al., 2018). The use of nanobodies in the future may also further improve the labeling of axonal proteins (Figure 3) and perhaps transcription factors. A further limitation of our protocol is the reliance on immersion fixed tissue sections for ECS preservation. While we focused on section thicknesses up to 1 mm, an important future goal is to achieve ECS preservation by perfusion of a whole brain (Cragg, 1980) and uniform permeabilization-free antibody labeling throughout large tissue volumes.

To replicate the protocol, we emphasize that it is important to first confirm that ECS preservation during acute section immersion fixation was successful, ideally by inspection of EM images. The degree of ECS preservation varies by brain region as previously reported (Pallotto et al., 2015) and needs to be calibrated for specific brain regions. The protocol also requires the typical optimization of IHC conditions for a given antibody such as concentration, buffer composition, and duration (Table 1). We have focused on proteins with known expression patterns for the optimization of the protocol. Our strategy was to compare the labeling pattern in permeabilized tissue to that of ECS preserved non-permeabilized tissue. For novel proteins, labeling specificity should be assayed as previously described (Lorincz and Nusser, 2008). We have not yet assessed whether antigen retrieval steps with proteases such as pepsin would further improve labeling while still maintaining ultrastructure (Watanabe et al., 1998) or whether multiplexing by repeated elution and re-labeling, such as in array tomography (Micheva and Smith, 2007), is compatible with the protocol.

The ultimate test of the utility of this protocol is in 3D correlative LM/EM reconstructions, for which we demonstrated two proof-of-principle applications (Figure 6). We anticipate that permeabilization-free pre-embedding IHC could play an important role in adding functional information to anatomical datasets. The combination of cellular-resolution functional imaging, permeabilization-free IHC and *en bloc*-based EM all within the same tissue would enable the correlation of function, cell-type identity, neuronal morphology and synaptic connectivity within local circuits. Our protocol could also be further optimized for tissue from species in which fluorescent genetic tagging of proteins is not possible, including human tissue obtained from immersion fixed biopsies.

## Materials and Methods

### Tissue fixation

C57BL/6J male mice (age p30 – p45) were used for all experiments except for the use of Pcdh21-Cre mice (Gensat stock # 030952-UCD; (Nagai et al., 2005)), age p30, to assess antibody labeling of mitral and tufted cells in the mouse olfactory bulb (*Figure 6*). All animal procedures were conducted in accordance with US National Institutes of Health guidelines, as approved by the National Institute of Neurological Disorders and Stroke Animal Care and Use Committee (ASP 1340).

For transcardially perfused tissue fixation, we followed a rapid perfusion approach (Tao-Cheng et al., 2007). Briefly, mice were anesthetized with isofluorane (Forane), and then perfused with 4% paraformaldehyde (PFA, Electron Microscopy Services) and 0.005% glutaraldehyde (GA, Electron Microscopy Sciences) in 175 mM sodium phosphate buffer (PB, pH 7.4). The brain was extracted and post-fixed for 4 hrs in the same fixative solution. The perfused brain was subsequently rinsed in 175 mM PB for 4 hrs and 300 μm coronal sections were cut on a Leica Vibratome.

For acute ECS preserved sections, mice were first anesthetized with isofluorane before swift decapitation. The brain was carefully removed from the skull, and 300 μm (up to 1 mm) coronal sections from the olfactory bulb, cerebral cortex, and/or hippocampus were cut on a vibratome (Leica) according to the procedure of (Bischofberger et al., 2006) and briefly stored in a cold carboxygenated (95% O_2_/5% CO_2_) ACSF solution (300-320 mOsm) containing (in mM): 124 NaCl, 3 KCl, 1.3 MgSO_4_.7H_2_O, 26 NaHCO_3_, 1.25 NaH_2_PO_4_.H_2_O, 20 Glucose, 2 CaCl_2_.2H_2_O. Sections were then immersion fixed using a protocol to preserve the extracellular space (Pallotto et al., 2015). The fixative concentrations were varied (see *Suppl Fig 2*) and an optimal primary fixative of 4% PFA + 0.005% GA in 175 mM PB (pH 7.4, 4 °C, 360 mOsm) for 4 hrs was chosen. The total duration between decapitation and fixation was less than 10 min.

### Immunohistochemistry

Primary and secondary antibody catalog numbers and incubation concentrations are listed in Supplemental Table 1. Optimal antibody concentrations were titrated for each antibody tested. For antibodies with stock concentrations of 0.5 mg/mL, antibody dilutions were typically 1:100 or 1:50. Antibody concentrations in nM were calculated using an IgG molecular weight of 150 kg/mol, the stock concentration and the dilution factor. All antibodies were diluted in an antibody buffer solution (ABS) of 170 mM NaCl + 10 mM PB (360 mOsm). 0.3% Triton X-100 (Electron Microscopy Sciences) was added to both the primary and secondary antibody solution for sections that were permeabilized. Sections were added to individual wells of a 96-well plate, each containing 50 - 75 μL ABS. Care was taken to ensure that the sections were fully immersed in the ABS and not adhered to the walls prior to wrapping the plate in parafilm. All IHC steps were carried out in the dark at room temperature on an orbital shaker at medium-high speed. The duration of primary and secondary antibody incubation was dependent on section thickness. Typically 300 μm sections were incubated in primary antibody for 72 hrs and in secondary antibody for 48 hrs.

#### Somatic labeling (300 μm thick sections)

Following fixation, sections were rinsed in 175 mM PB for 4 hrs, then a buffered glycine solution (50 mM glycine in 175 mM PB) for 16 hrs, and then rinsed for 6 hrs in 175 mM PB. Regions of 300 μm thick coronal sections were then trimmed down to volumes of approximately 2 × 2 × 0.3 mm^3^ with a scalpel for cortical and hippocampal regions; olfactory bulb sections were left intact. For labelling of NeuN-positive somata (*Figures 1, 5a,b,e,f*), sections were incubated in the ABS with Alexa Fluor-488 conjugated anti-NeuN (Abcam) for 72 hrs. For labelling of calbindin-positive and calretinin-positive somata (*Figures 3a-b, 5c,d,g,h*), sections were simultaneously incubated in guinea pig anti-calbindin (Synaptic Systems) and rabbit anti-calretinin (Sigma) for 72 hrs, rinsed in 175 mM PB for 12 hrs, and then incubated in the secondary antibodies, DyLight-405 goat anti-guinea pig IgG F(ab’)_2_ fragment (Jackson ImmunoResearch) for calbindin and Alexa Fluor-594 donkey anti-rabbit IgG F(ab’)_2_ fragment (Jackson ImmunoResearch) for calretinin, for 48 hrs.

#### Somatic labeling (1 mm thick sections)

Coronal sections were fixed in 4% PFA + 0.005% GA in either 100 mM, 175 mM, or 225 mM PB (pH 7.4, 4 °C) for 4 hrs. Following fixation, sections were rinsed for 4 hrs in PB of the respective concentration used for fixation, then a buffered glycine solution (50 mM glycine also in the respective PB concentration) for 16 hrs, and then rinsed for 6 hrs in PB of the respective concentration. 1 mm thick coronal sections were trimmed down to volumes of approximately 4 × 1 × 1 mm^3^ with a scalpel and incubated in Alexa Fluor-488 conjugated anti-NeuN (Abcam) for 120 hrs (*Figure 2*). Sections were then embedded in 10% quick dissolve agarose (GeneMate) and hemisected to a volume of approximately 2 × 1 × 1 mm^3^ to observe anti-NeuN penetration through the 1 mm depth with 2P microscopy on the hemisected surface.

#### Axon labeling

Following fixation, sections were rinsed in 175 mM PB for 4 hrs, then a buffered glycine solution (50 mM glycine in 175 mM PB) for 16hrs, and then rinsed for 6 hrs in 175 mM PB. Regions of 300 μm thick coronal sections were then trimmed down to volumes of approximately 2 × 2 × 0.3 mm^3^ with a scalpel. For labelling of choline acetyl transferase-positive axons (ChAT, *Figure 3c-d*), sections were incubated in goat anti-ChAT (EMD Millipore) for 72 hrs, rinsed in 175 mM PB for 8 hrs, and then incubated in DyLight-594 donkey anti-goat (Abcam) for 48 hrs. For labelling of tyrosine hydroxylase-positive axons (TH, *Figure 6d*), sections were incubated in Alexa Fluor-488 conjugated anti-tyrosine hydroxylase, clone LNC1 (EMD Millipore) for 72 hrs, and then rinsed in 175 mM PB for 8 hrs.

#### Synaptic protein labeling

Following fixation, sections were rinsed in 175 mM PB for 4 hrs, then a buffered glycine solution (50 mM glycine in 175 mM PB) for 16hrs, and then rinsed for 6 hrs in 175 mM PB. Regions of 300 μm thick coronal sections were then trimmed down to volumes of approximately 2 × 2 × 0.3 mm^3^ with a scalpel. For labelling of Homer in postsynaptic terminals (*Figure 4a-b*), sections were incubated in rabbit anti-Homer 1 (Synaptic Systems) for 72 hrs, rinsed in 175 mM PB for 8 hrs, and then incubated in Alexa Fluor-594 donkey anti-rabbit IgG F(ab’)_2_ fragment (Jackson ImmunoResearch) for 48 hrs. For labelling of synaptophysin in presynaptic terminals (*Figure 4c*), sections were first incubated in Alexa Fluor-488 conjugated rabbit anti-synaptophysin (Abcam) for 72 hrs, rinsed in 175 mM PB for 24 hrs, and then incubated in Alexa Fluor-594 conjugated rabbit anti-synaptophysin (Abcam) plus 0.3% Triton for 48 hrs.

#### Serum blocking

For sections in which serum blocking was employed (*Suppl Fig 4)*, tissue was first incubated in a blocking buffer of 10% normal donkey serum, 1% bovine serum albumin and 0.05% sodium azide in 175 mM PB for 4 hrs. Following the serum blocking step, sections were incubated with antibodies as described above with 3% normal donkey serum.

#### HRP labeling of eGFP

Cre-dependent expression of eGFP in axons and dendrites of mitral cells was achieved by injection of adeno-associated virus encoding eGFP into the anterior piriform cortex. AAV pCAG-FLEX-EGFP-WPRE was a gift from Hongkui Zeng (Addgene plasmid # 51502-AAV1 ; http://n2t.net/addgene:51502 ; RRID:Addgene_51502) (Oh et al., 2014). Under isofluorane anesthesia, age p30 Pcdh21-Cre mice were positioned into a stereotaxic head holder, and a small craniotomy was made on the dorsal surface above the injection site. A glass pipette was used to inject the virus, targeting axon terminals of mitral cells bilaterally in the anterior piriform cortex (350 nL per side). After 3 weeks animals were sacrificed and 300 μm thick acute ECS preserved coronal sections of the olfactory bulb were prepared as above. Following fixation, sections were rinsed in 175 mM PB for 4 hrs, then a buffered glycine solution (50 mM glycine in 175 mM PB) for 16hrs, and finally rinsed for 6 hrs in 175 mM PB. eGFP fluorescence in the coronal sections was first imaged with a 2P microscope without SeeDB clearing (see below). Subsequently, cytosolic eGFP in mitral and tufted cells was labeled for EM (*Figure 6a-c*) by incubating sections in rabbit anti-GFP (ThermoFisher) for 72 hrs, rinsing in 175 mM PB for 8 hrs, and then incubating in horseradish peroxidase (HRP)-conjugated donkey anti-rabbit (Abcam) for 48 hrs. Sections were re-fixed with 2% GA in 150 mM PB for 2 hrs at room temperature, rinsed in 150 mM PB for 2 hrs, rinsed in glycine (50 mM glycine in 175 mM PB) for 16 hrs, rinsed again in 150 mM PB for 2 hrs, and then incubated at room temperature in the dark on a rotator in 1.4 mM diaminobenzidine hydrotetrachloride (DAB, Serva) and 0.56 mM hydrogen peroxide (Sigma) in 175 mM PB for 10 hrs to polymerize DAB in the presence of HRP.

### Tissue clearing

Following antibody labeling, sections were rinsed in 175mM PB for 16 hrs, re-fixed in buffered PFA (2% PFA in 175 mM PB) for 2 hrs, rinsed in 175 mM PB for 4 hrs. Sections were subsequently incubated in a modified SeeDBp protocol (Ke et al., 2013): 20% w/v fructose in 175 mM PB for 2 hrs, 40% w/v fructose in 175 mM PB for 3 hrs, 60% w/v fructose in 175 mM PB for 4 hrs, 80% w/v fructose in 175 mM PB for 4-6 hrs, 100% w/v fructose in water for a minimum of 10 hrs, and finally in SeeDB solution (80.2% w/w fructose in water and 0.5% 1-thioglycerol) for a minimum of 10 hrs. Fructose incubation was omitted for 1 mm thick sections because the hemisected surface was imaged directly to assess antibody penetration.

### Two-photon imaging

Immunolabeled sections were mounted in SeeDB solution between two #1 cover slips in a custom imaging chamber and imaged with a 20x, 1.0 NA water immersion objective (Olympus) on a 2P laser scanning microscope (Sutter) using ScanImage (Pologruto et al., 2003). An excitation wavelength of 770 - 800 nm was used to image sections incubated with Alexa Fluor-488 and Alexa Fluor-594/DyLight-594. An excitation wavelength of 770 nm was used to image the DyLight-405 fluorophore. 2P image stacks were collected with 1 μm z-step sizes throughout the depth of 300 μm thick sections at various x-y pixel sizes.

### Near-infrared branding

A near-infrared branding technique (Bishop et al., 2011) was used to burn fiducial marks 20 - 30 μm below the top surface of sections around a region of interest (*Figure 6e*) after 2P image stack acquisition, typically a 150 × 150 μm^2^ region. Following branding, sections were gradually returned to 175 mM PB in decreasing fructose concentrations over 16 hrs and then post-fixed in 2% GA in 150 mM sodium cacodylate buffer (CB, Electron Microscopy Sciences) for 2 hrs.

**Figure 6.**
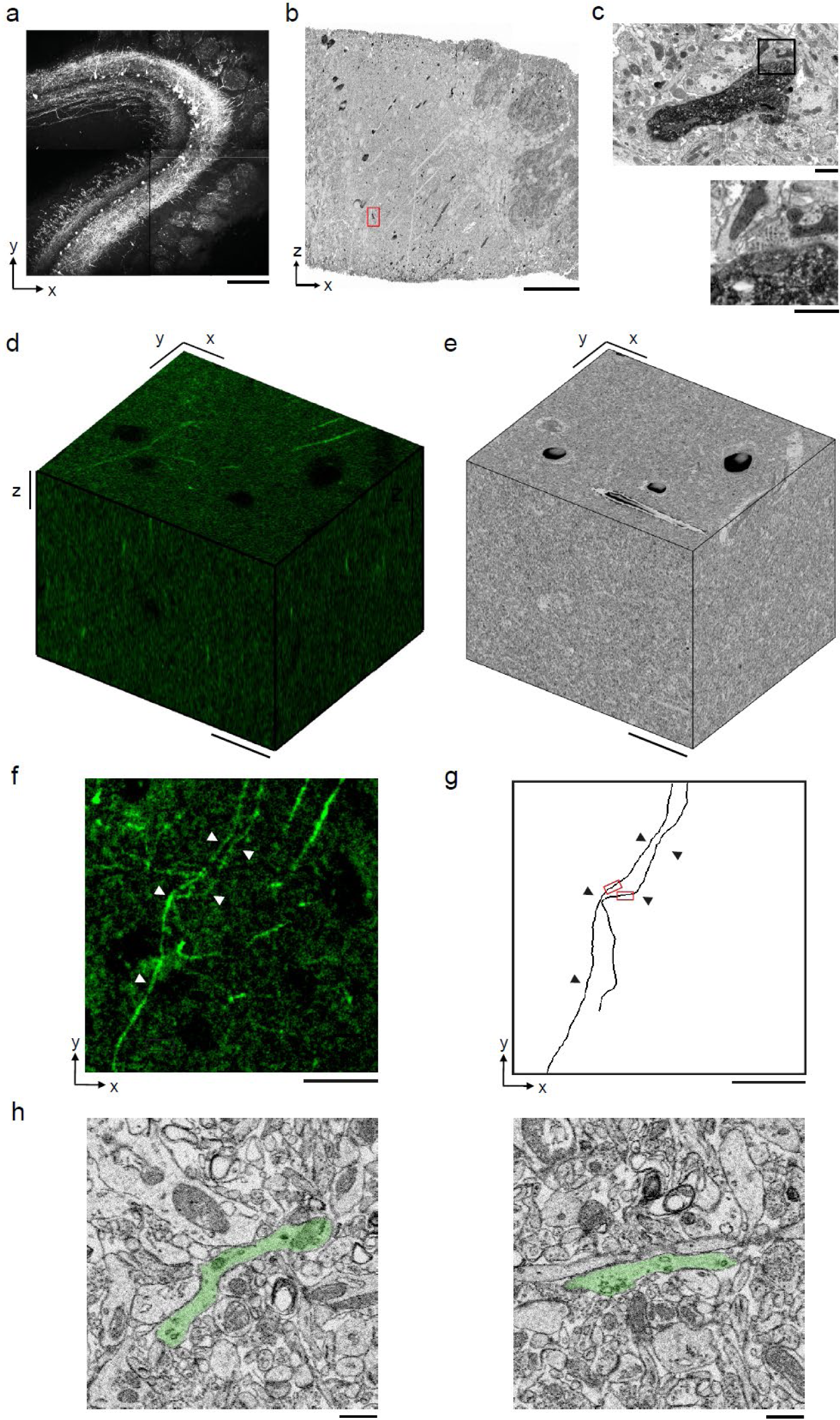
Permeabilization-free labeling for correlative microscopy. **a-c)** Correlative microscopy of mitral cells (MCs) in the mouse olfactory bulb. **a)** 300 μm thick acute ECS preserved sections containing MCs expressing eGFP following AAV transfection. **b)** Incubation with anti-GFP and HRP-conjugated secondary antibody was performed without Triton, followed by polymerization of DAB and EM staining. **c)** Enlarged regions of panel b (red rectangle) demonstrating confinement of DAB product to MC dendrites (upper panel) and a presynaptic terminal formed on a labeled dendrite (lower panel). **d-h)** Correlative microscopy of TH+ axons in the mouse medial prefrontal cortex. **d)** 2P image stack from a 300 μm thick ECS preserved acute section that was incubated with anti-tyrosine hydroxylase (TH) without Triton to label dopaminergic axons. **e)** A SBEM volume centered on branded fiducial marks. **f)** A 2 μm thick X-Y average intensity projection from panel d with TH+ axons indicated (white arrowheads). **g)** Anatomical reconstructions of corresponding TH+ axons from the SBEM volume with matching axons indicated (black arrowheads). **h)** Example EM images of the reconstructed axons from the regions indicated in panel g. Axons are false-colored (green). Scale bars: a: 200 μm; b: 50 μm; c: 1 μm; d-g: 20 μm; h: 1 μm.

### Tissue processing for electron microscopy

For sections prepared for correlative EM, sections were thoroughly rinsed in 175 mM CB prior to EM staining. Sections were stained using a modification of previously described protocols (Briggman et al., 2011; Karnovsky, 1971). Briefly, tissue was stained in a solution containing 2% osmium tetroxide, 3% potassium ferrocyanide, and 2mM CaCl_2_ in 150 mM CB for 2 hrs at 4 °C. The osmium stain was amplified with 1% thiocarbohydrazide (1 hr at 50 °C), and 2% osmium tetroxide (1 hr at room temperature). The tissue was then stained with 1% aqueous uranyl acetate for 6 hrs at 45 °C and lead aspartate for 6 hrs at 45 °C. The tissue was dehydrated at 4 °C through an ethanol series (70%, 90%, 100%), transferred to propylene oxide, infiltrated at room temperature with 50%/50% propylene oxide/Epon Hard, and then 100% Epon Hard (Glauert and Lewis, 1998). The blocks were cured at 60 °C for 24 h. 50 – 70 nm ultrathin sections were cut from block faces and imaged on a NanoSEM 450 (FEI).

### Serial block-face electron microscopy

For SBEM acquisition (*Figure 6e*), data were collected using a custom serial block-face microtome designed by K.L. Briggman. The specimens were cut out of the flat-embedding blocks and re-embedded in Epon Hard on aluminum stubs. The samples were then trimmed to a block face of ∼200 μm wide and ∼200 μm long. The samples were imaged in a scanning electron microscope with a field-emission cathode (NanoSEM 450, FEI). Back-scattered electrons were detected using a concentric segmented back-scatter detector. The incident electron beam had an energy of 2.4 keV and a current of ∼200 pA. Images were acquired with a pixel dwell time of 2 us and size of 10.25 nm × 10.25 nm. Imaging was performed at high vacuum, with the sides of the block sputter coated with a 100 nm thick layer of gold. The section thickness was set to 32 nm. 2354 consecutive block faces were imaged from the sample, resulting in aligned data volume of 8704 × 8064 × 2354 voxels (corresponding to an approximate spatial volume of 89 × 83 × 75 um^3^).

Following SBEM acquisition, we aligned the 2P image stack to the SBEM volume with an affine transformation using soma and blood vessel locations as landmarks. Axons in the SBEM volume corresponding to TH+ axons in the 2P stack were annotated using Knossos (https://knossos.app/) (Helmstaedter et al., 2011).

### Co-localization analysis

For the colocalization analysis of synaptophysin (*Figure 4d*), we used a method similar to a previously described rotation-based strategy (Soto et al., 2011). Two simultaneously acquired fluorescent 2P stacks represented the labeling of synaptophysin without Triton (AF488 green channel) and subsequently with Triton (AF594 red channel). We analyzed whether presynaptic puncta were labeled with Triton permeabilization (AF594, red channel) that had not been previously labeled without Triton permeabilization (AF488, green channel). Fifty presynaptic puncta were manually outlined with 0.4 μm diameter ROIs and the voxel intensities averaged within each ROI in the (AF594) red channel. These ROI intensities were plotted against the same ROI intensities in the (AF488) green channel to assess the co-localization of the synaptophysin labeling before and after Triton permeabilization. We then compared this correlation to a 90 degree rotation of the (AF488) green channel. A linear regression t-test was used to test the statistical significance of the co-localized correlation compared to the rotated correlation.

## Acknowledgements

We thank B. Fubara for assistance with animal perfusions and M. Pallotto for assistance with virus injections. We also thank J. Diamond, S. Haverkamp and M. Pallotto for comments of the manuscript.

---

**Suppl Figure 1.**
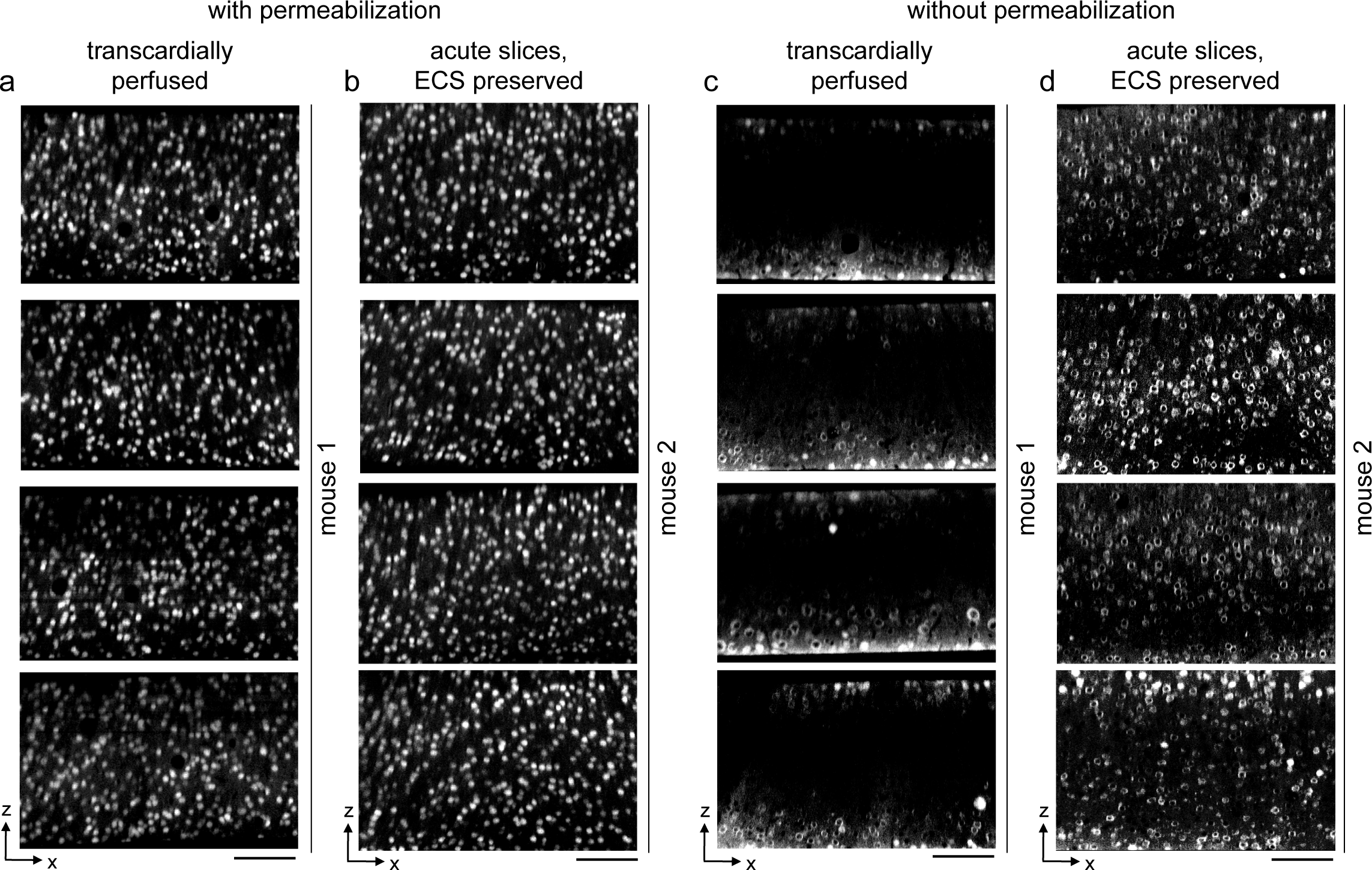
Replication of permeabilization-free labeling across sections and animals. **a,b)** 300 μm thick sections from a transcardially perfused mouse (a, n = 4 sections) and a mouse from which acute ECS preserved sections were collected (b, n = 4 sections). Sections were labeled with Alexa Fluor-488 conjugated anti-NeuN with 0.3% Triton. 2P image volumes were acquired. X-Z reslices are 10 μm average intensity projections from the center of the sections. **c,d)** 300 μm thick sections from a transcardially perfused mouse (c, n = 4 sections) and a mouse from which acute ECS preserved sections were collected (d, n = 4 sections). Sections were labeled with Alexa Fluor-488 conjugated anti-NeuN without permeabilization. 2P images as in panels a,b. Scale bars: 100 μm.

**Suppl Figure 2.**
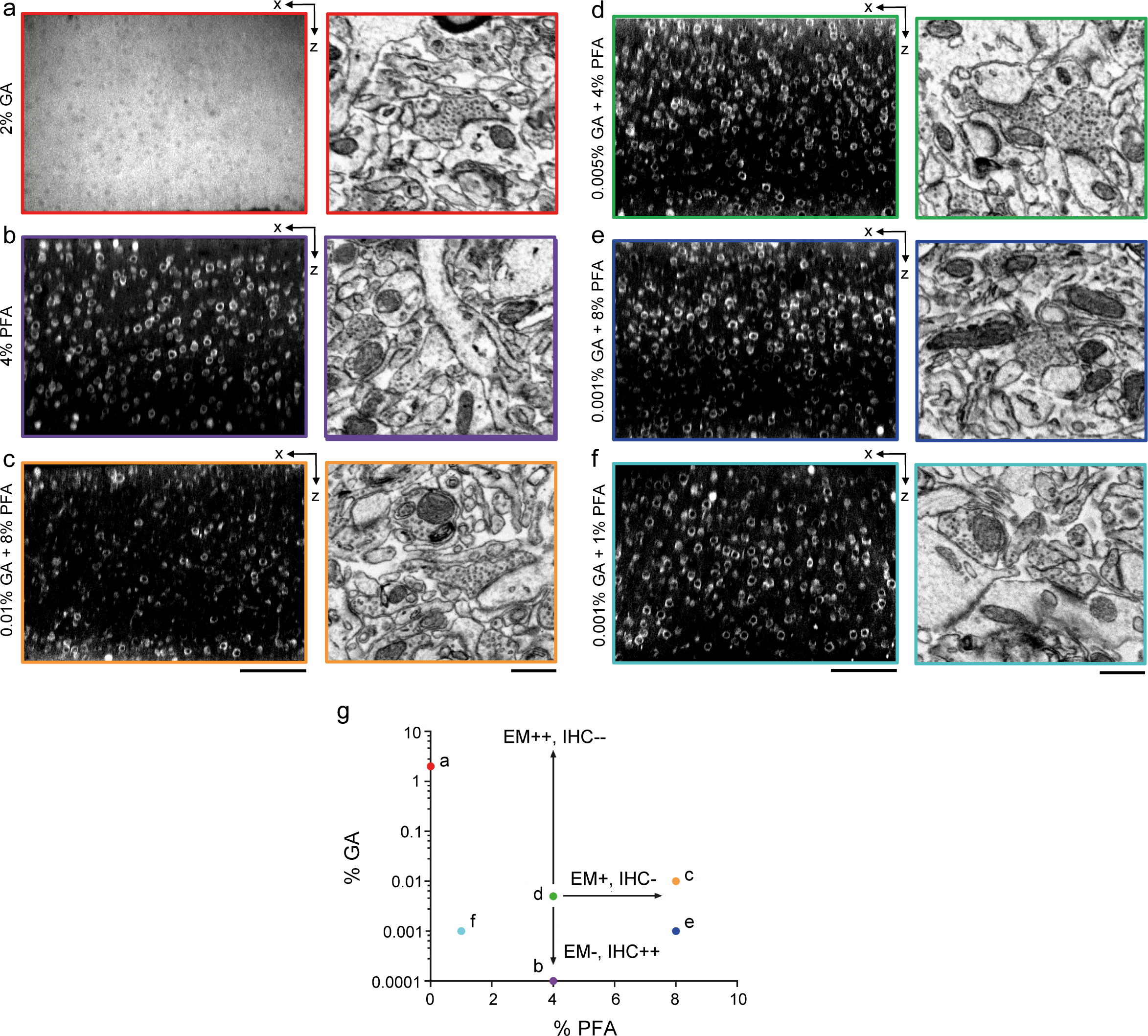
Optimization of fixation parameters. **a-f)** 300 μm thick acute ECS preserved sections from mouse cerebral cortex. Slices were fixed with different concentrations of glutaraldehyde (GA) and paraformaldehyde (PFA) and then incubated with Alexa Fluor-488 conjugated anti-NeuN. 2P image volumes were acquired. X-Z reslices (left panels) are 10 μm average intensity projections from the center of the sections. Sections were then fixed with 2% GA, stained for EM and a region from the center of the section was examined for ultrastructural integrity (right panels). **g)** The points in parameter space that were tested are color-coded to match the panels in a-f. A combination of 0.005% GA + 4% PFA (panel d) was selected as an optimal tradeoff between ultrastructural quality and antibody penetration. Scale bars: a: 100 μm; b-d: 100 μm (upper panels), 0.5 μm (lower panels).

**Suppl Figure 3.**
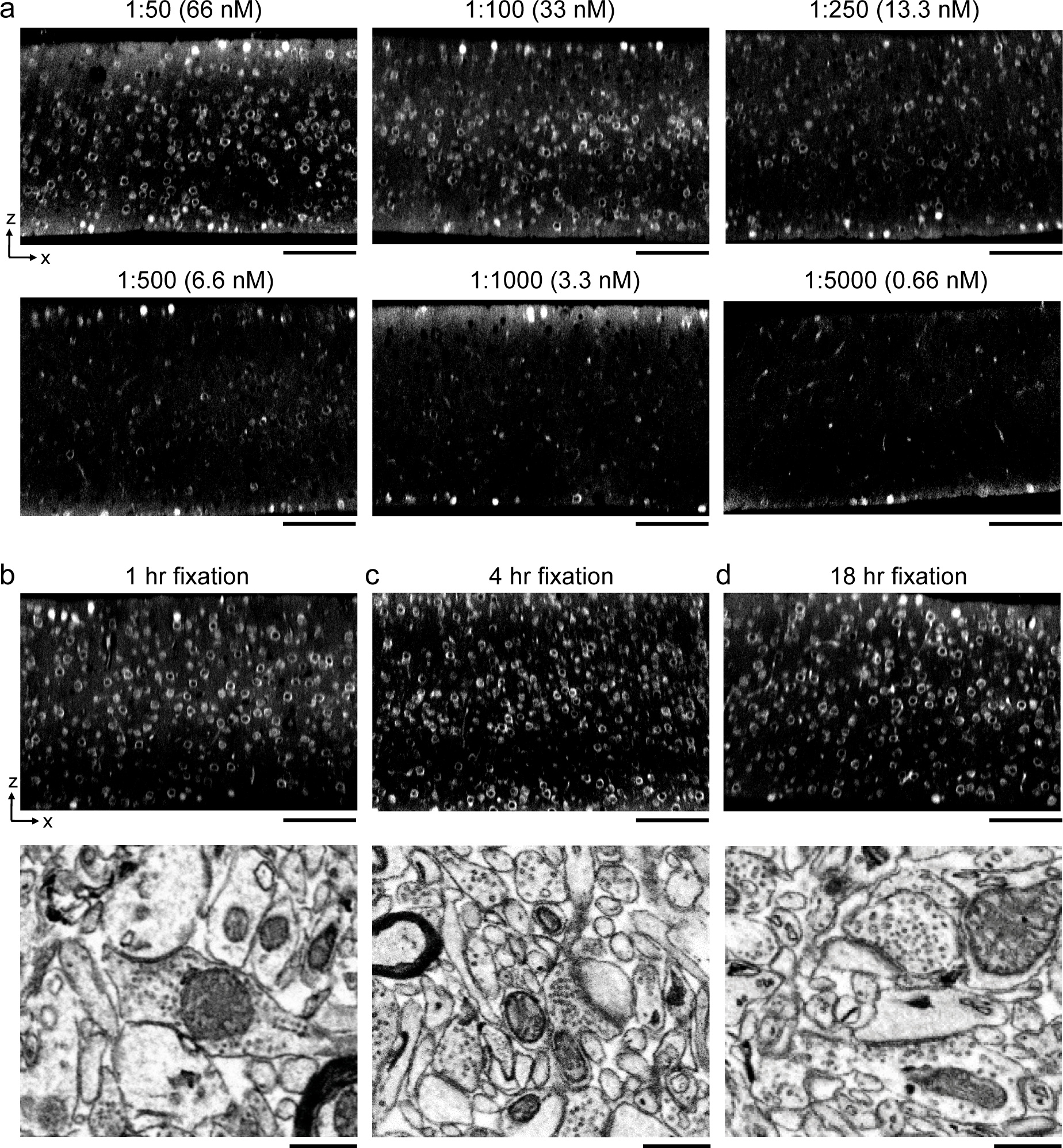
Optimal antibody concentration and duration of primary fixation. **a)** 300 μm thick acute ECS preserved sections from mouse cerebral cortex. Sections were incubated in different concentrations of Alexa Fluor-488 conjugated anti-NeuN as indicated for 72 hrs. 2P image volumes were acquired. X-Z reslices are 10 μm average intensity projections from the center of the sections. **b-d)** 300 μm thick acute ECS preserved sections from mouse cerebral cortex. Sections were fixed for varying durations (1, 4, or 18 hrs) with 0.005% GA + 4% PFA prior to incubation with Alexa Fluor-488 conjugated anti-NeuN. 2P image volumes were acquired. X-Z reslices (upper panels) are 10 μm average intensity projections from the center of the sections. Sections were then stained for EM and a region from the center of the section was examined for ultrastructural integrity (lower panels). Scale bars; upper panels: 100 μm; lower panels: 0.5 μm.

**Suppl Figure 4.**
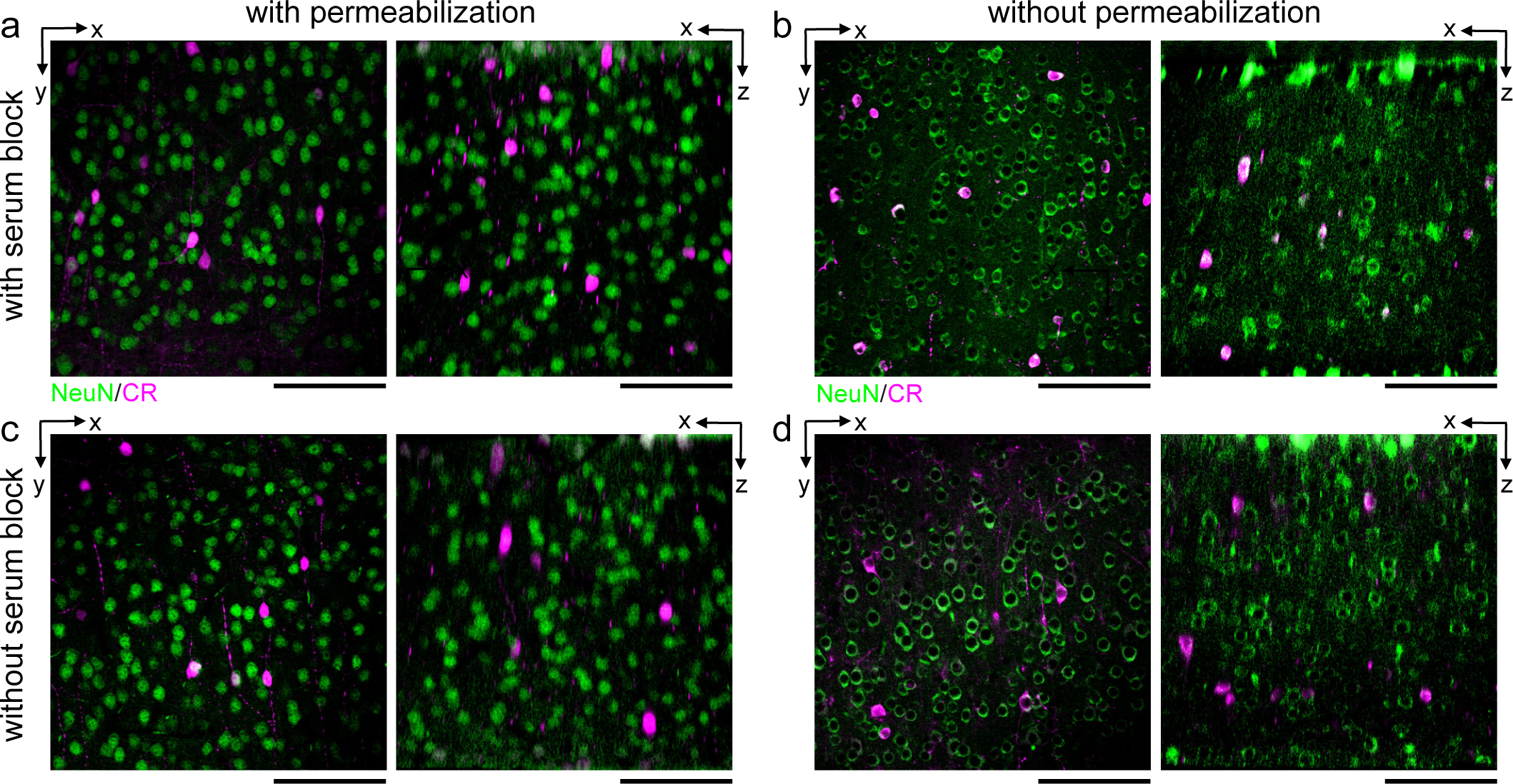
Comparison of the effect of serum blocking on labeling thick sections. 300 μm thick acute ECS preserved sections from mouse cerebral cortex. Sections were incubated with Alexa Fluor-488 conjugated anti-NeuN and anti-calretinin (CR) either with (a,b) or without (c,d) 3% donkey serum and with (a,c) or without (b,d) permeabilization with 0.3% Triton. 2P image volumes were acquired. X-Y (left panels) and X-Z reslices (right panels) are 10 μm average intensity projections from the center of the sections. Scale bars: 100 μm.

**Suppl Figure 5.**
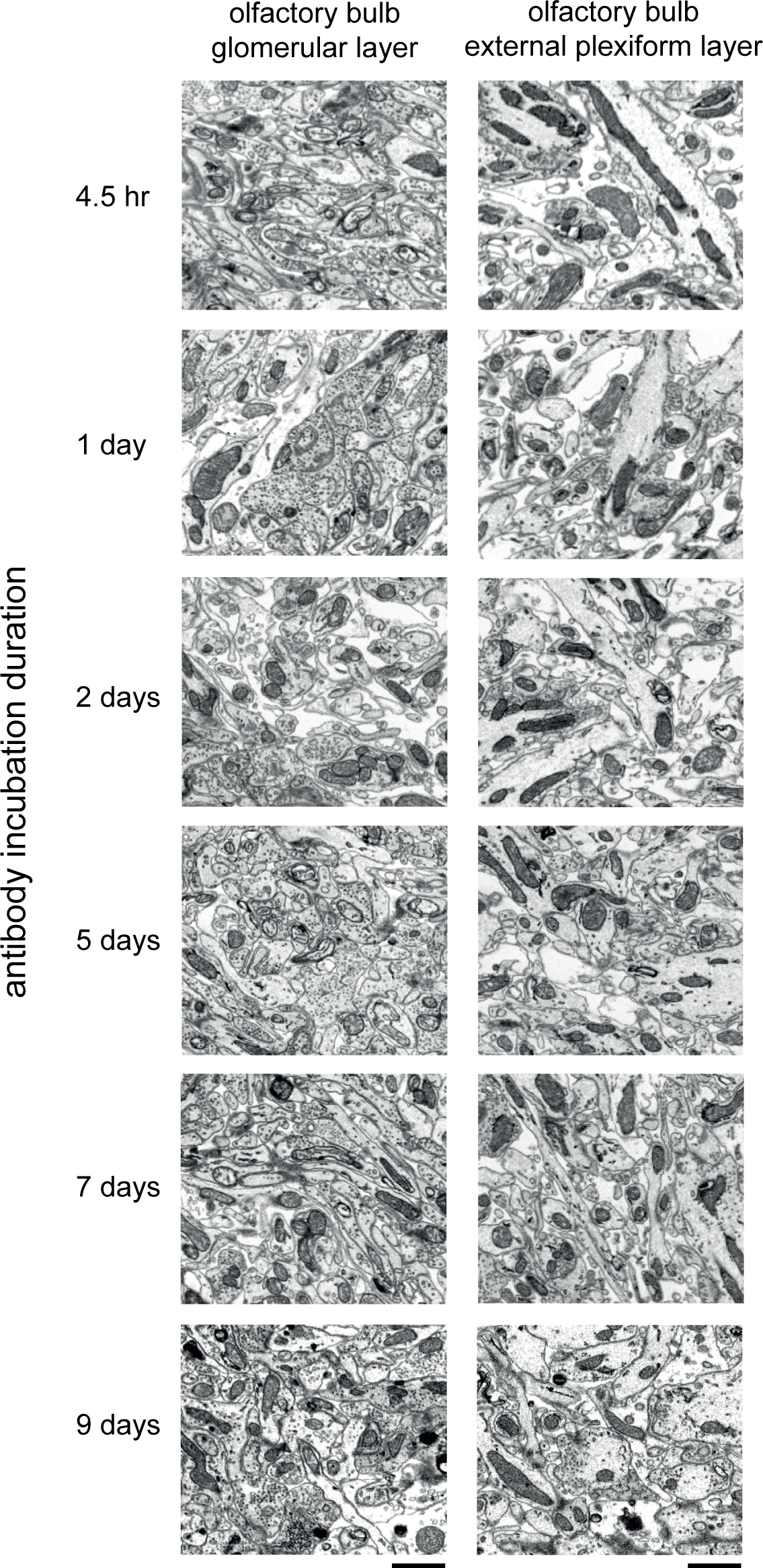
Comparison of the effect of prolonged antibody incubation on ultrastructure quality. 300 μm thick acute ECS preserved sections from mouse olfactory bulb. Sections were incubated with Alexa Fluor-488 conjugated anti-NeuN for varying durations (4.5 hrs to 9 days). Sections were then stained for EM and a region from the center of the section was examined for ultrastructural integrity in the glomerular layer (left panels) and external plexiform layer (right panels). Scale bars: 1 μm.

**Suppl Figure 6.**
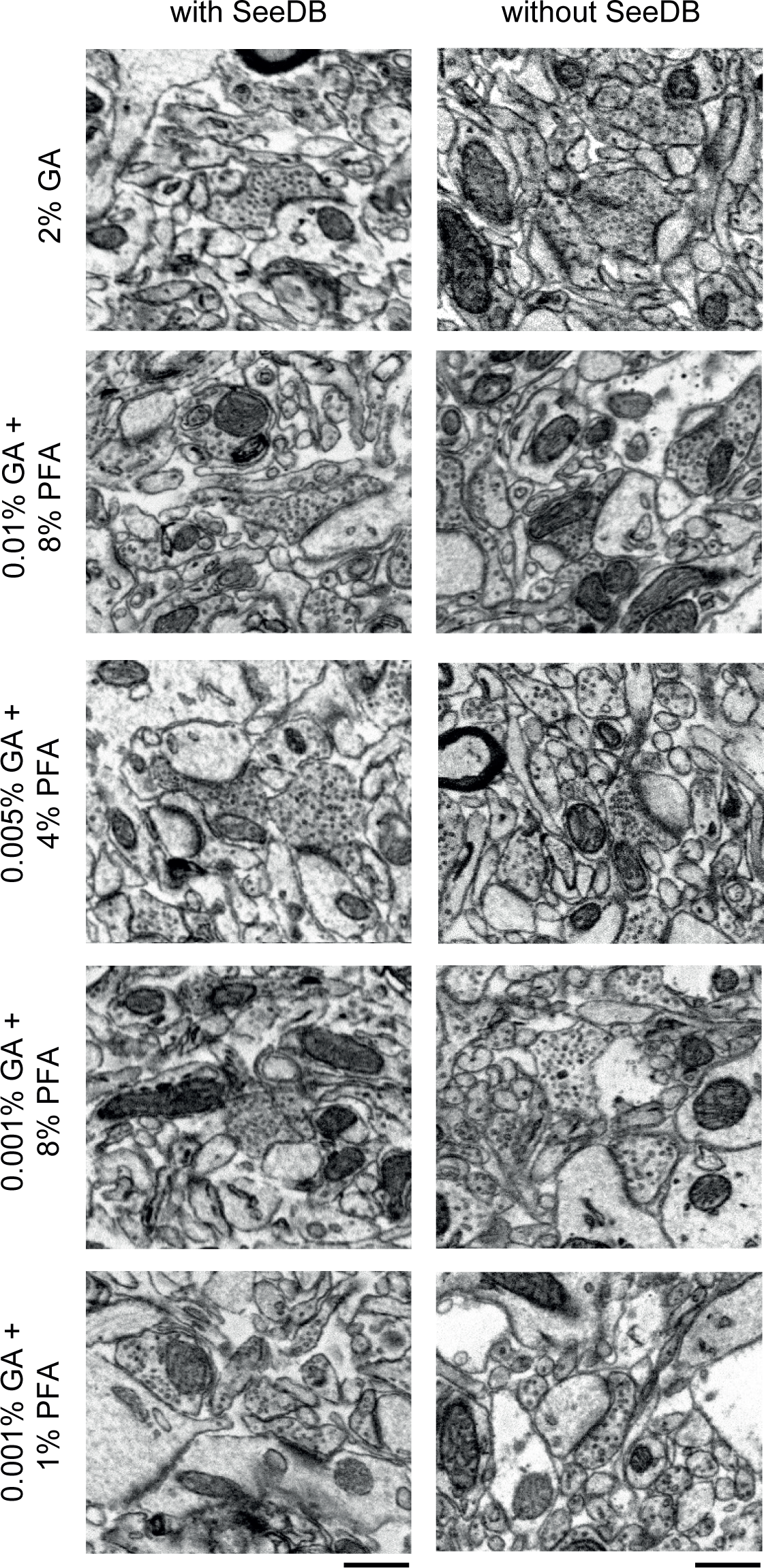
Comparison of the effect of SeeDB clearing on ultrastructure quality. 300 μm thick acute ECS preserved slices from mouse cerebral cortex. Slices were fixed with different concentrations of GA and PFA as indicated and were then incubated with Alexa Fluor-488 conjugated anti-NeuN. Half of the slices then underwent the SeeDB procedure (see Methods) and the others remained in buffer. The SeeDB slices were then rehydrated to buffer and then all sections were refixed in 2% GA and stained for EM. Scale bars: 0.5 μm.

**Suppl Table 1.** Table of tested primary and secondary antibodies. Primary and secondary antibodies that successfully labeled protein targets throughout the depth of 300 μm thick acute ECS preserved brain slices.

| Primary Antibody | Company and Cat # | Stock Concentration | Concentration | Target |
| --- | --- | --- | --- | --- |
| Anti-calbindin (CB) | Synaptic Systems<br>214 005 | 1 mg/mL | 1:100 | Calcium binding protein |
| Anti-calretinin (CR) | Sigma<br>C7479 | 0.8 mg/mL | 1:50 | Calcium binding protein |
| Anti-tyrosine hydroxylase (TH) | EMD Millipore<br>MAB318-AF488 | 0.5 mg/mL | 1:100 | Enzyme |
| Anti-choline acetyltransferase (ChAt) | EMD Millipore<br>AB144P | 0.5 mg/mL | 1:100 | Enzyme |
| Anti-synaptophysin (Syn) (Alexa Fluor-594) | Abcam<br>ab206868 | 0.5 mg/mL | 1:50 | Presynaptic protein |
| Anti-synaptophysin (Syn) (Alexa Fluor-488) | Abcam<br>ab196379 | 0.5 mg/mL | 1:50 | Presynaptic protein |
| Anti-Homer 1 | Synaptic Systems<br>160 003 | 1 mg/mL | 1:50 | Postsynaptic protein |
| Anti-NeuN | Abcam<br>ab190195 | 0.5 mg/mL | 1:50 | Neuronal marker |
| Anti-GFP | ThermoFisher<br>G10362 | 0.2 mg/mL | 1:50 | Marker protein |

| Secondary Antibody | Company and Cat # | Stock Concentration | Concentration |
| --- | --- | --- | --- |
| Alexa Fluor 594 F(ab')<sub>2</sub> fragment donkey anti-rabbit | Jackson ImmunoResearch<br>711-586-152 | 1.5 mg/mL | 1:100 |
| DyLight 594 donkey anti-goat | Abcam<br>ab96937 | 0.5 mg/mL | 1:100 |
| DyLight 405 goat anti-guinea pig F(ab')<sub>2</sub> fragment specific | Jackson ImmunoResearch<br>106-475-006 | 1.5 mg/mL | 1:25 |
| HRP Donkey anti-rabbit | Abcam<br>ab205722 | 2 mg/mL | 1:50 |

## Notes

### Competing Interest Statement

The authors have declared no competing interest.

